# Senescence during early differentiation reduced chondrogenic differentiation capacity of mesenchymal progenitor cells

**DOI:** 10.1101/2023.03.09.531873

**Authors:** Chantal Voskamp, Wendy J. L. M. Koevoet, Gerjo J.V.M. van Osch, Roberto Narcisi

## Abstract

Mesenchymal stromal/progenitor cells (MSCs) are promising for cartilage cell-based therapies due to their differentiation capacity. However, MSCs can become senescent during *in vitro* expansion, a state characterized by stable cell cycle arrest, metabolic alterations, and substantial changes in the gene expression and secretory profile of the cell. In this study, we aimed to investigate how senescence and the senescence associated secretory phenotype (SASP) affect chondrogenic differentiation of MSCs. To study the effect of senescence, we exposed MSCs to gamma irradiation during expansion or during chondrogenic differentiation (pellet culture). When senescence was induced during expansion or at day-7 of chondrogenic differentiation, we observed a significant reduction in cartilage matrix. Interestingly, when senescence was induced at day-14 of differentiation, chondrogenesis was not significantly altered. Moreover, exposing chondrogenic pellets to medium conditioned by senescent pellets had no significant effect on the expression of anabolic or catabolic cartilage markers, suggesting a neglectable paracrine effect of senescence on cartilage generation in our model. Finally, we show that senescent MSCs had lower phosphorylated SMAD2 levels after TGFβ1stimulation compared to control MSCs. Overall, these results suggest that the occurrence of senescence in MSCs during expansion or early differentiation could be detrimental for cartilage tissue engineering.

## Introduction

Articular cartilage is prone to damage and has a limited repair capability. Full-thickness loss of articular cartilage does not regenerate spontaneous and can lead to the degenerative joint disease osteoarthritis (OA) ^1, 2^. Current treatment methods such as microfracture or autologous chondrocyte graft implantation have limitations and fail to prevent OA progression ^3^. An alternative strategy to repair damaged cartilage uses mesenchymal stem/stromal cells (MSCs). MSCs are progenitor cells that can be isolated from several tissues such as bone marrow, synovial membrane and adipose tissue and have the capacity to differentiate towards the chondrogenic lineage ^4, 5^. To obtain enough MSCs to repair a cartilage defect, *in vitro* expansion is necessary. During extensive expansion, MSCs gradually lose their chondrogenic differentiation capacity ^6, 7^, limiting the applications of these cells. Expansion also triggers cellular senescence, a process leading to an irreversible cell cycle arrest, major metabolic changes and a senescence-associated secretory phenotype (SASP) ^8, 9^. SASP factors produced by senescent cells include IL-6, IL-8, IL-1β, TNFα, MMP3 and MMP13 ^10-12^. It is known that these SASP factors can hamper tissue regeneration ^13^, for example exposure to TNFα and IL-1β during *in vitro* chondrogenesis limit the chondrogenic differentiation capacity of MSCs ^14^. In addition, SASP factors such as TNFα and IL-1β are known pro-inflammatory factors contributing to the pathophysiology of OA ^15, 16^. This is further supported by the fact that transplantation of senescent fibroblasts can lead to an OA-like phenotype, including cartilage erosion and delamination of the articular surface ^17^. In addition, the SASP factors such as CCL2, IL-6, IGFBP4 and IGFBP7 have been suggested to contribute to the spread of cellular senescence in MSC ^18, 19^, known as paracrine senescence ^20^. It is known that cellular senescence alters the differentiation capacity of MSCs, especially the effects on the osteogenic and adipogenic lineages are studied. Loss of osteogenic and adipogenic potential has been demonstrated in senescent MSC ^6^, however it has also been reported that in late passaged MSCs the levels of mineralized matrix declines, while adipocytes differentiation increase ^21, 22^, indicating the complexity of this phenomena. Moreover, cartilage displays a decline in repair capacity with aging ^23^, but little is known about the effect of cellular senescence on the chondrogenic differentiation capacity of MSCs. The aim of this study was therefore to determine how cellular senescence and their SASP affect chondrogenesis of MSCs.

## Method

### MSC isolation and expansion

Iliac crest bone chips were obtained from patients (9-13 years) undergoing alveolar bone graft surgery N=13. The tissue was procured as leftover/waste surgical material and it was reviewed and deemed exempt from full ethical review after ethical approval by the Erasmus Medical Ethical Committee (MEC-2014-16,). These pediatric MSCs have been previously characterized and used in this study because they exhibit a low number of senescent cells at early passages ^18, 24^. MSCs were isolated by rinsing bone chips twice with 10 mL alpha-MEM (Gibco brand ThermoFisher Scientific, Waltham, MA, USA) supplemented with 10% fetal calf serum (brand ThermoFisher Scientific; selected batch 41Q2047K), 1.5 μg/ml fungizone (Invitrogen brand ThermoFisher Scientific), 50 μg/ml gentamicin (Invitrogen brand ThermoFisher Scientific), 1 ng/ml FGF2 (R&D Systems, Minneapolis, MN, USA) and 0.1 mM ascorbic acid-2-phosphate (Sigma-Aldrich, Zwijndrecht, the Netherlands). The MSCs were plated in T175 flasks and after 24 hours the non-adherent cells were washed away. MSCs were trypsinized at sub-confluency and reseeded in a density of 2,300 cells/cm^2^. MSCs between passage 3 and 6 were used for experiments.

### Irradiation of MSCs in monolayer followed by chondrogenic differentiation

Senescence was induced in the cells using 20 Gy ionizing radiation by a RS320 X-Ray machine (X-Strahl, Camberley, UK) ^25^. MSCs in monolayer were irradiated in a T175 flask (60-70% confluency) for 22 min (20 Gy). 24 hours post-irradiation the cells were trypsinized and seeded at a 9,600 cell/cm^2^ density. Mock irradiated MSC were used as non-senescent controls and seeded at 2,300 cells/cm^2^. 7 days post-irradiation, irradiated and non-irradiated MSCs were trypsinized, mixed (0, 25, 50, 75 and 100% irradiated versus non-irradiated cells) and centrifuged at 300 x g for 8 min to obtain pellets of 2×10^5^ cells. To induce chondrogenesis, cell pellets were cultured in chondrogenic medium, containing DMEM-HG medium (Invitrogen brand ThermoFisher Scientific), supplemented by 1% ITS (BD, Franklin Lakes, NJ, USA), 1.5 μg/ml fungizone (Invitrogen brand ThermoFisher Scientific), 50 μg/ml gentamicin (Invitrogen brand ThermoFisher Scientific), 1 mM sodium pyruvate (Invitrogen brand ThermoFisher Scientific), 40 μg/ml proline (Sigma-Aldrich), 10 ng/ml TGFβ1 (R&D Systems), 0.1 mM ascorbic acid-2-phosphate (Sigma-Aldrich) and 100 nM dexamethasone (Sigma-Aldrich) for 7, 14 or 21 days. The medium was renewed twice a week.

### Senescence-associated beta-galactosidase staining

To confirm cellular senescence, 7 days post-irradiation, cells from each donor (N=5) were trypsinized and seeded in monolayer cultures in triplicates. Subconfluent cells were washed twice with PBS. Next, the cells were fixed with 0.5% glutaraldehyde and 1% formalin in Milli-Q water for 5 min at room temperature. Then the cells were washed twice with Milli-Q water and subsequently the cells were stained with freshly made X-gal solution containing 0.5% X-gal, 5 mM potassium ferricyanide, 5 mM potassium ferrocyanide, 2mM MgCl2, 150mM NaCl, 7mM C6H8O7 and 25mM Na2HPO4 incubated for 24 hours at 37°C. Cells were counterstained with 1:25 pararosaniline detected with bright field microscopy. Two independent researchers scored at least 100 cells as negative or positive as previously described ^25^.

### Irradiation of chondrogenic pellets and conditioned medium

To induce cellular senescence in chondrogenic pellets. Non-irradiated MSCs were cultured in chondrogenic medium and renewed twice a week. MSCs in pellets were irradiated at day 7 or 14 of chondrogenic differentiation in a 15 mL tube for 22 min (20 Gy). Chondrogenic medium was renewed 24 hours after irradiation, next the medium was renewed twice a week. Mock irradiated cells/pellets were used as controls.

To determine the effect of SASP factors on chondrogenic differentiation, we generated two different sets of chondrogenic pellets from the same donor, medium donating pellets from irradiated MSCs and medium recipient pellets from non-irradiated MSCs. To determine the effect at different time points during chondrogenesis we analyzed the RNA expression of the medium recipient pellets at day 9 and at day 16. First, to generated conditioned medium, the medium of the donating pellets was replaced by DMEM-HG medium supplemented with 1% ITS, 1.5 μg/ml fungizone (Invitrogen brand ThermoFisher Scientific), 50 μg/ml gentamicin (Invitrogen brand ThermoFisher Scientific), 1 mM sodium pyruvate (Invitrogen brand ThermoFisher Scientific) and 40 μg/ml proline 24-48 hours before harvesting. The medium from the donating pellets (N=2) was collected and pooled per donor and time point. To remove cell debris, the medium was centrifuged at 14,000 g for 1 min. Next, medium was mixed with DMEM-HG medium supplemented with 1% ITS, 1.5 μg/ml fungizone (Invitrogen brand ThermoFisher Scientific), 50 μg/ml gentamicin (Invitrogen brand ThermoFisher Scientific), 1 mM sodium pyruvate (Invitrogen brand ThermoFisher Scientific) and 40 μg/ml proline at ratio 3:1, and 0.1 mM ascorbic acid-2-phosphate (Sigma-Aldrich) and 10 ng/ml TGFβ1 was added to the total volume. The conditioned medium mixture was added to non-irradiated recipient MSCs pellets for 2 consecutive days, specifically at day 7- and 8 (timepoint 9 days), or at day 14- and 15 (timepoint 16 days) during chondrogenic differentiation. 24 h after the last addition of conditioned medium, at day 9 and day 16 respectively, the medium recipient pellets were lysed in RNA-STAT (Tel-Test, Friendswood, TX, USA) for mRNA expression analysis. Media from non-irradiated medium donating MSC pellets using cells from the same donor and at the same time points, were generated and used as a control conditioned media.

### (Immuno)Histochemistry chondrogenic pellets

Pellets were fixed with 3.7% formaldehyde after 7, 14 or 21 days of chondrogenic induction. Next, pellets were embedded in paraffin and sectioned at 6 μm. To detect glycosaminoglycans, sections were stained with 0.04% thionine solution. To detect collagen type-2, sections were first treated with 0.1% Pronase (Sigma-Aldrich) in PBS for 30 min at 37°C, followed by 1% hyaluronidase (Sigma-Aldrich) in PBS for 30 min at 37°C. Sections were incubated with 10% normal goat serum (Sigma-Aldrich) and 1% bovine serum albumin (Sigma-Aldrich) in PBS for 30 min, followed by incubation with the collagen type-2 antibody (II-II 6B3, Developmental Studies Hybridoma Bank) for 1h. Then samples were incubated with a biotin-conjugated antibody (HK-325-UM, Biogenex) for 30 min, followed by incubation with alkaline phosphatase-conjugated streptavidin (HK-321-UK, Biogenex) for 30 min. New Fuchsin chromogen (B467, Chroma Gesellschaft) was used as a substrate. As a negative control an IgG1 isotype antibody (X0931, Dako Cytomation) was used. The positive area per pellet was determined using ImageJ software.

### Osteogenic and adipogenic differentiation

To induce osteogenic differentiation, expanded MSCs were trypsinized, seeded at a density of 1.2×10^4^ cells/cm^2^ and cultured in DMEM HG medium (Gibco brand ThermoFisher Scientific) with 10% fetal calf serum (Gibco brand ThermoFisher Scientific), 1.5 μg/ml fungizone (Invitrogen brand ThermoFisher Scientific), 50 μg/ml gentamicin (Invitrogen brand ThermoFisher Scientific), 10 mM β-glycerophosphate (Sigma-Aldrich), 0.1 μM dexamethasone (Sigma-Aldrich) and 0.1 mM ascorbic acid-2-phosphate (Sigma-Aldrich) for 12-21 days. To detect calcium deposits the cultures were fixed in 3.7 % formaldehyde, followed by hydration with Milli-Q water and incubation with 5% silver nitrate solution (Von Kossa; Sigma Aldrich) for 1 h in the presence of bright light. Next the cultures were washed with distilled water followed by counterstaining with 0.4% thionine (Sigma-Aldrich). MSCs were used in triplicates (N=3 donors). To induce adipogenic differentiation, expanded MSCs were trypsinized, seeded in a density of 2×10^4^ cells/cm^2^ and cultured in DMEM HG (Gibco brand ThermoFisher Scientific) with 10% fetal calf serum (Gibco brand ThermoFisher Scientific), 1.5 μg/ml fungizone (Invitrogen brand ThermoFisher Scientific), 50 μg/ml gentamicin (Invitrogen brand ThermoFisher Scientific), 1.0 μM dexamethasone (Sigma-Aldrich), 0.2 mM indomethacin (Sigma-Aldrich), 0.01 mg/ml insulin (Sigma-Aldrich) and 0.5 mM 3-isobutyl-l-methyl-xanthine (Sigma-Aldrich) for 21 days. To detect intracellular lipid accumulation, cells were fixed in 3.7 formaldehyde, followed by incubation with 0.3% Oil red O solution (Sigma-Aldrich) for 10 min and washes with distilled water. MSCs were used in triplicates (N=3 donors).

### DNA and Glycosaminoglycan (GAG) Quantification

Pellets were digested at day 21 of chondrogenic differentiation using 1 mg/ml Proteinase K, 1 mM iodoacetamide, 10 μg/ml Pepstatin A in 50 mM Tris, 1 mM EDTA buffer (pH 7.6; all Sigma-Aldrich) for 16 h at 56°C, followed by Proteinase K inactivation at 100°C for 10 min. Afterwards, to determine the amount of DNA, cell lysates were treated with 0.415 IU heparin and 1.25 μg RNase for 30 min at 37°C followed by addition of 30 μl CYQUANT GR solution (Invitrogen). Samples were analyzed using a SpectraMax Gemini plate reader with an excitation of 480 nm and an emission of 520 nm. As a standard, DNA sodium salt from calf thymus (Sigma-Aldrich) was used. To determine the amount of GAG, cell lysates were incubated with 1,9-dimethylmethylene blue (DMB) as previously described by Ferndale et al. ^26^, and analyzed with an extinction of 590 nm and 530 nm. The 530:590 nm ratio was used to determine the glycosaminoglycan concentration. As a standard chondroitin sulfate sodium salt from shark cartilage (Sigma-Aldrich) was used.

### mRNA Expression analysis

For both MSCs in pellet cultures and MSCs in monolayer cultures, the medium was renewed 24 hours before cell lysis. Pellets were washed twice with PBS, lysed in RNA-STAT (Tel-Test) and manually homogenized. Next, RNA was isolated using chloroform and purified using the RNeasy micro kit (Qiagen, Hilden, Germany) following the manufacturer’s protocol. MSCs in monolayer were washed twice with PBS and RNA was isolated using RLT lysis buffer supplemented with 1% β-mercaptoethanol. Subsequently, RNA was purified using the RNAeasy micro kit using the manufacturer’s protocol. The RevertAid First Strand cDNA synthesis kit (Fermentas brand ThermoFisher Scientific) was used to reverse transcribe the RNA to cDNA. Next, real-time polymerase chain reactions were done with SYBR Green (Fermentas brand ThermoFisher Scientific) and TaqMan (Applied Biosystems brand ThermoFisher Scientific) MasterMix on a CFX96TM PCR machine (Bio-Rad, Hercules, CA, USA) using different primers listed in **Table 1**. Genes with a housekeeping function are often used as reference genes for qPCR analysis, however senescent cells often have altered their housekeeping functions ^27^. Therefore, we tested four different housekeeping genes (*GAPDH, HPRT1, RPS27A* and *ACTB*) for each dataset and only used the genes that were stable across the different conditions as reference. Gene expression levels were calculated using the 2^−ΔCt^ formula.

**Table 1.**
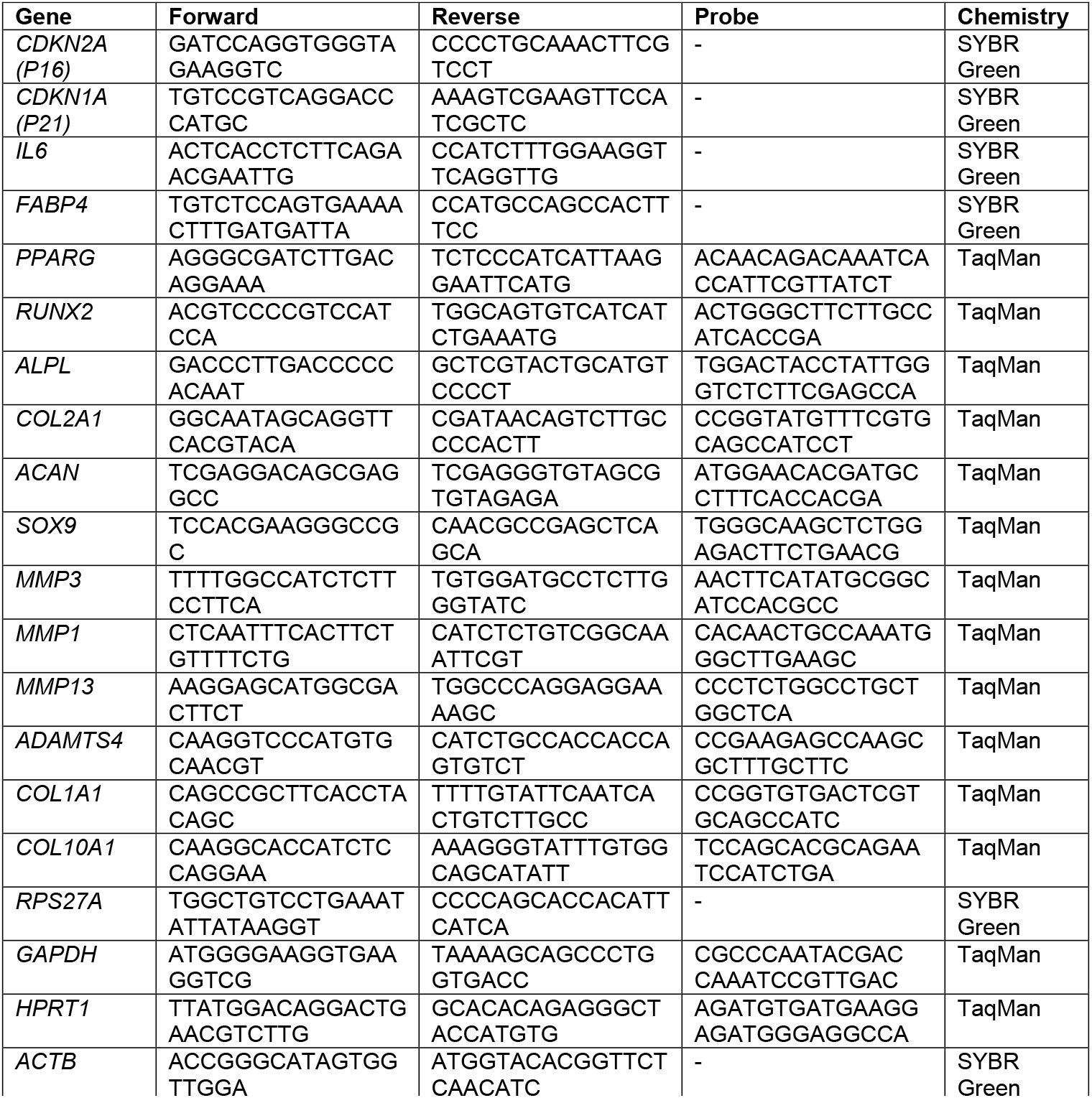
Primer sequences.

### Western blot

Irradiated MSCs and non-irradiated MSCs in monolayer were serum starved for 16 h in alpha-MEM (Invitrogen) supplemented with 1% BSA, 1.5 μg/ml fungizone (Invitrogen) and 50 μg/ml gentamicin (Invitrogen). Next, MSCs were stimulated with 0 or 10 ng/ml TGFβ1 for 30 min and subsequently, cells were lysed in MPER lysis buffer (ThermoFisher Scientific) with 1% Halt Protease Inhibitor (ThermoFisher Scientific) and 1% Halt Phosphatase Inhibitor (ThermoFisher Scientific). Protein samples, from MSCs from different donors (N=3 donors, in triplicates), were separated on a 4-12% SDS-PAGE gel (ThermoFisher Scientific) by electrophoresis using an equal amount of protein (5-12 μg) per sample. Proteins were transferred semi-wet from the SDS-PAGE gel on a nitrocellulose membrane (Millipore). The membrane was transferred to a 5% dry milk powder blocking solution in <u>T</u>ris-<u>B</u>uffered <u>S</u>aline with 0.1% <u>T</u>ween-20 (Millipore Sigma; TBST) for 3 h. Next, the membrane was incubated with the primary monoclonal rabbit antibody against phospho-SMAD2 Ser465/Ser467 (Cell Signaling technology; 3108S;) using a 1:1000 dilution in 5% BSA in TBST overnight at 4°C. Later, the membrane was incubated with a secondary anti-rabbit antibody conjugated with peroxidase (Cell Signaling, 7074S) using a 1:1000 dilution in 5% dry milk powder in TBST for 1.5 h at room temperature. Phospho-SMAD2 signal was detected with the SuperSignal Wester Pico Complete Rabbit IgG detection kit (ThermoFisher Scientific).

### Data analysis

The Kolmogorov-Smirnov test was used to verify the normal (Gaussian) distribution of all the histology, RNA expression and western blot data. For statistical evaluation, a linear mixed model was applied, using the different conditions as fixed parameters and the donors as random factors. Bonferroni post-hoc test was used to correct for multiple comparisons. Data analysis was performed using PSAW statistics 20 software (SPSS Inc., Chicago, IL, USA). *P*-values less than 0.05 were considered as statistically significant.

## Results

### Cellular senescence impaired the chondrogenic capacity of MSCs

Cellular senescence was induced in monolayer MSCs using gamma irradiation (20 Gy) and confirmed by an increased mRNA expression of cell-cycle dependent *CDKN2A* (6.9-fold) and *CDKN1A* (4.8-fold), a higher mRNA expression of the SASP associated gene *IL6* (8.6-fold) and a higher percentage of senescence associated β-galactosidase positive cells than the mock treated control MSCs (0 Gy; **Figure 1A-B**). After 21 days of chondrogenic induction, irradiated MSCs, had an impaired capacity to deposit the typical chondrogenic extracellular proteins GAG and COL2 (**Figure 1C-D**). To determine whether senescent MSCs have an overall reduced differentiation capacity or whether it was specific for the chondrogenic lineage, we assessed their osteogenic and adipogenic differentiation capacity. After adipogenic differentiation, the cells show lipid accumulation and expression of adipogenic genes *PPRG* and *FABP4* in both the irradiated and non-irradiated cells, (**Figure 1E-F**) although for *FABP4* a reduced expression was detected compared to control MSCs.. After osteogenic differentiation, irradiated and non-irradiated cells show no significant differences in the osteogenic markers *RUNX2* and *ALPL* (**Figure 1G-H**). Overall, these results indicate that senescent MSCs can differentiate towards the adipogenic and the osteogenic lineage, while a strong negative effect was detected specifically for the chondrogenic differentiation.

**Figure 1.**
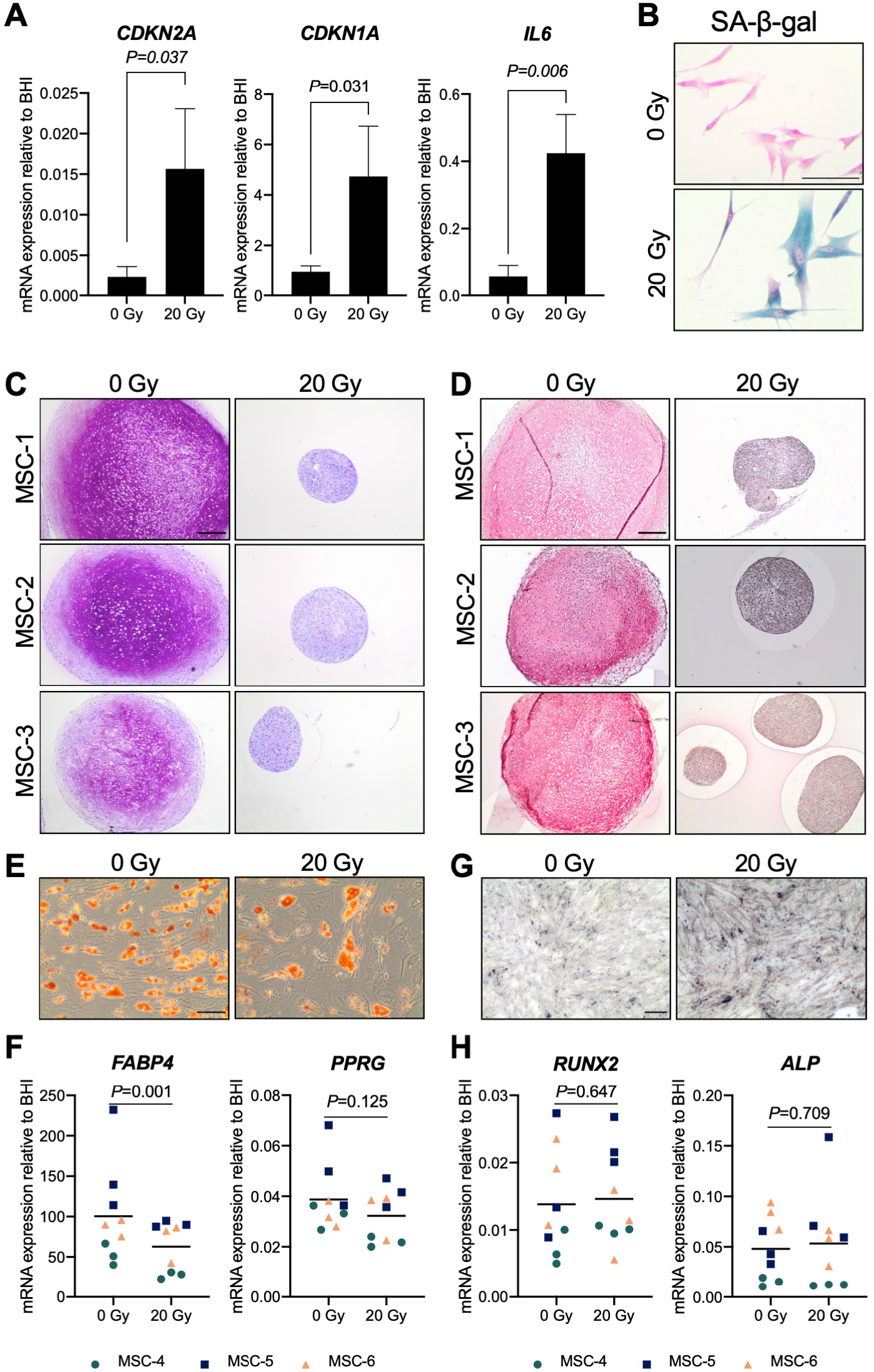
Chondrogenic differentiation was impaired in senescent MSCs. MSCs that were gamma irradiated (20 Gy) or mock irradiated (0 Gy) after expansion. (A) *CDKN2A (P16*), *CDKN1A* (P21) and *IL6* mRNA relative to the best housekeeping index (BHI; *GAPDH, HPRT, RPS27A* and *ACTB*). N=3 donors with 2-3 replicates per donor. Data show grand mean and standard deviation. (B) Representative images of MSCs stained for senescence-associated β-galactosidase (SA-β-gal) activity. Scale bar represents 100 μm. N=3 donors with 2-3 replicates per donor. (C-D) Representative images of Thionine (C) and Collagen type 2 (D) staining of pellets of (mock-)irradiated MSCs that were chondrogenically differentiated for 21 days. Scale bar represents 250 μm. N=3 donors with 3 pellets per donor. (E) Representative images of Oil red O staining of (mock-)irradiated MSCs that were differentiated towards adipogenic lineage for 21 days. Scale bar represents 100 μm, N=3 donors with 3 replicates per donor. (F) *FABP4* and *PPARG* mRNA expression relative to the best housekeeping index (BHI; *GAPDH, RPS27A* and *ACTB*) of MSCs that were differentiated towards adipogenic lineage for 21 days. N=3 donors with 3 replicates per donor. (G) Representative images of Von Kossa staining of (mock-)irradiated MSC that were differentiated towards osteogenic lineage for 14-21 days. Scale bar represents 200 μm, N=3 donors with 3 replicates per donor. (H) *RUNX2* and *ALP* mRNA expression relative to the best housekeeping index (BHI; *GAPDH, RPS27A* and *ACTB*) of MSCs that were differentiated towards osteogenic lineage for 14-21 days. N=3 donors with 3 replicates per donor. Data show individual data points and grand mean. *P*-values were obtained with the linear mixed model, using the different irradiation conditions as fixed parameters and the donors as random factors.

### Senescence during early MSC differentiation inhibited cartilage formation

In order to understand whether cellular senescence is affecting chondrogenic differentiation only when induced in specific differentiation stages, we used non senescent MSCs to generate pellets and triggered senescence by irradiation during chondrogenic differentiation. Specifically, we induced senescence in pellet cultures by gamma irradiation (20 Gy) at 7 or 14 days of chondrogenesis, in a 21-day differentiation protocol. As expected, mock treated pellets (0 Gy) had an increased GAG deposition over time and the deposition is highest at day 21 of chondrogenic differentiation (*P*=0.028 compared to day 7), while pellets treated with 20 Gy at day 7 of culture had an average of 1.6-fold reduction of GAG deposition at day 21 compared to controls (**Figure 2A** and **Supplementary Figure 1**; *P*=0.035). Immunostaining revealed an overall similar pattern between COL2 and GAG deposition, with a lower COL2 deposition detected at day 21 in day7-irradiated pellets compared to control pellets (**Figure 2B** and **Supplementary Figure 2**; *P*=0.010). At gene expression level, *COL2A1* and *ACAN* significantly increased over time in both irradiated and control conditions, but at day 21 the day7-irradiated pellets showed a significant reduced expression compared to control (**Figure 2C;** *COL2A1* and *ACAN*). The transcription factor *SOX9* did not strongly increase over time and its expression was lower in day7-irradiated pellets compared to control at day 21 (**Figure 2C;** *SOX9*). Between day 14 and day 21 of chondrogenic differentiation, gene expression of *COL2A1, ACAN* and *SOX9* remained similar (*P*=1.000).

**Figure 2.**
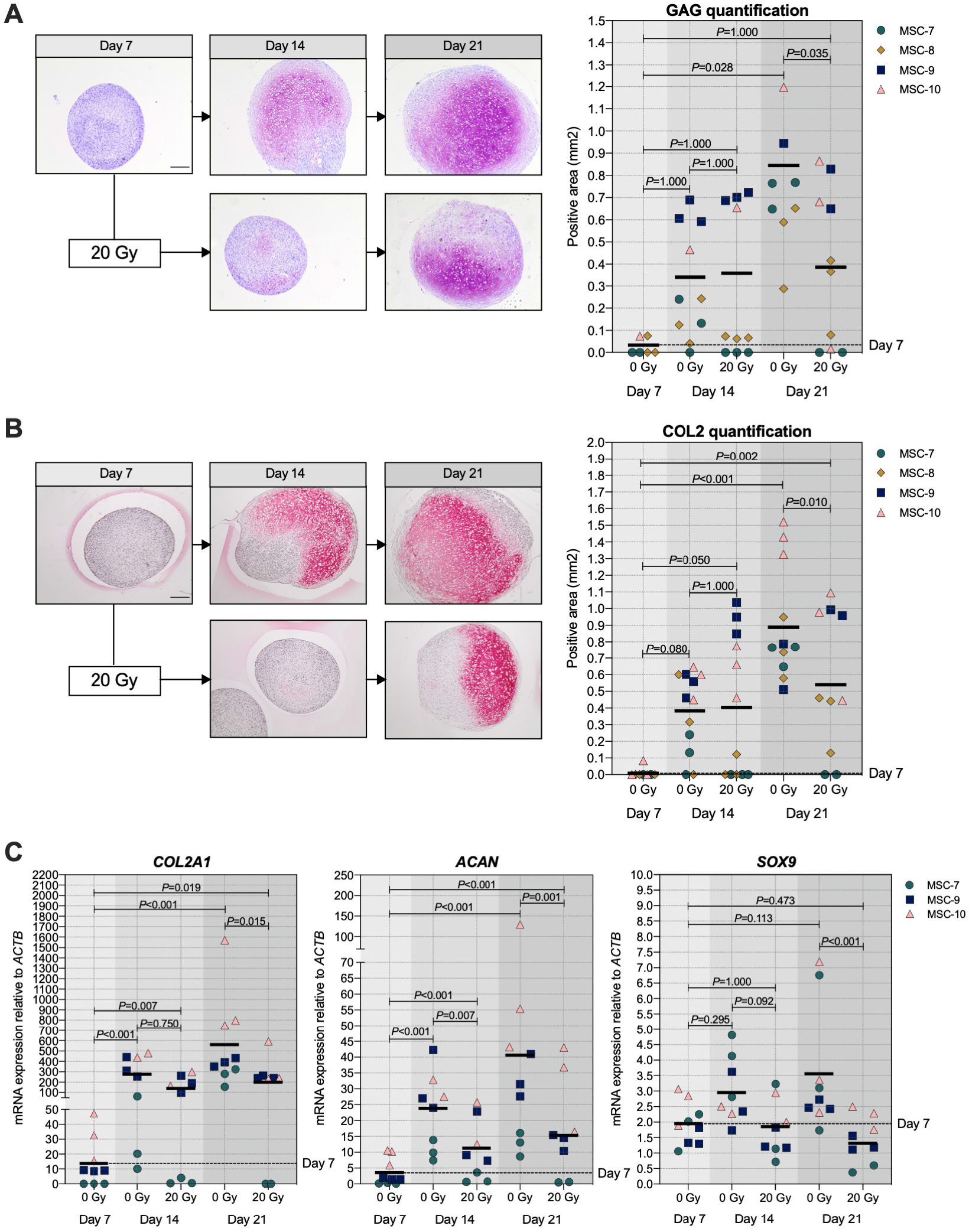
20 Gy irradiation at day 7 during MSC differentiation reduced chondrogenic markers at day 21. (A-B; left panels) Representative images of (A) Thionine (GAG) and (B) Collagen type 2 (COL2) staining of MSC control pellets that were chondrogenically differentiated for 7, 14 and 21 days or MSC pellets that were irradiated at day 7 during chondrogenic differentiation and subsequently differentiated for 7 or 14 days. The scale bar represents 200 μm. (A-B; right panels) Quantification of (A) GAG or (B) COL2 positive area per condition in mm^2^. N=4 donors with 1-3 replicates per donor. (C) Gene expression of chondrogenic markers in MSC control pellets that were chondrogenically differentiated for 7, 14 and 21 days or MSC pellets that were irradiated at day 7 during chondrogenic differentiation and subsequently differentiated for 7 and 14 days. Gene expression levels were normalized using *ACTB*. N=3 donors with 2-3 replicates per donor. Data show individual data points and grand mean. *P*-values were obtained with the linear mixed model, using the different irradiation conditions as fixed parameters and the donors as random factors.

Interestingly, when we irradiated the pellets at day14 the deposition of GAG and COL2 did not change compared to non-irradiated controls (**Figure 3A-B** and **Supplementary Figure 3**). Similarly, *COL2A1, ACAN* and *SOX9* gene expression at day 21 were comparable between day-14 irradiated pellets and controls (**Figure 3C**). Overall, these data suggest that the chondrogenesis of MSCs was not negatively influenced by irradiation at day 14. To test whether there was at least an effect on the known hypertrophic tendency of MSCs during chondrogenesis, *COL10A1, ALPL* and *RUNX2* expression were analyzed. No differences in *COL10A1, ALPL* and *RUNX2* expression were observed between day14-irradiated and mock treated pellets (**Figure 3D**). Although with donor variation, these data suggest that senescence during early differentiation (day-7) inhibits chondrogenic maturation, while senescence during late chondrogenesis (day-14) has no evident effect.

**Figure 3.**
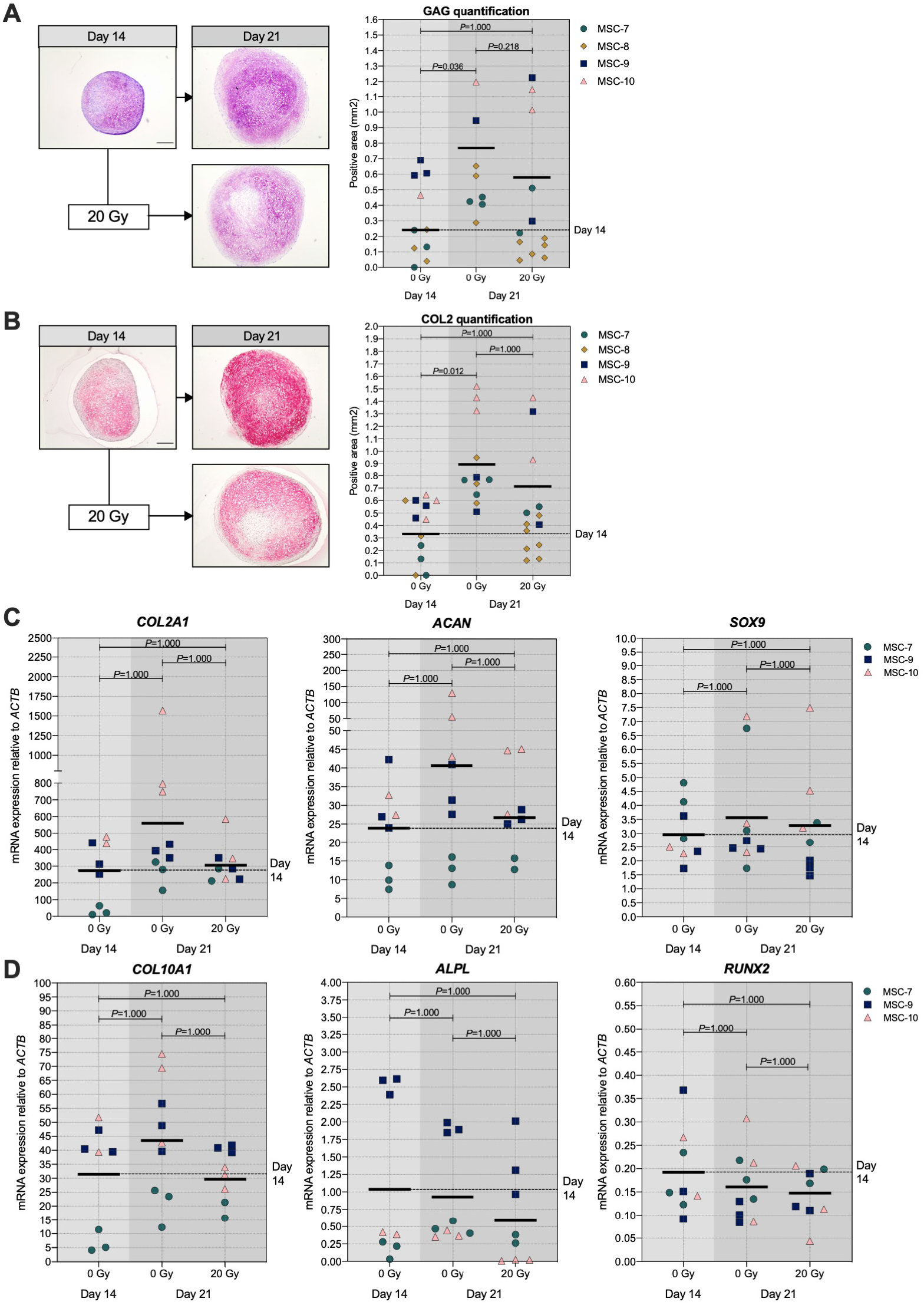
Irradiation at day 14 during MSC differentiation did not alter chondrogenic markers at day 21. (A-B; left panels) Representative images of (A) Thionine (GAG) and (B) Collagen type 2 (COL2) staining of MSC control pellets that were chondrogenically differentiated for 14 and 21 days or MSC pellets that were irradiated at day 14 during chondrogenic differentiation and subsequently differentiated for 7. The scale bar represents 200 μm. (A-B; right panels) Quantification of (A) GAG or (B) COL2 positive area per condition in mm^2^. N=4 donors with 1-7 replicates per donor. (C-D) Gene expression of (C) chondrogenic markers and (D) hypertrophic markers in MSC control pellets that were chondrogenically differentiated for 14 and 21 days or MSC pellets that were irradiated at day 14 during chondrogenic differentiation and subsequently differentiated for 7 days. Gene expression levels were normalized using *ACTB*. N=3 donors with 2-3 replicates per donor. Data show individual data points and grand mean. *P*-values were obtained with the linear mixed model, using the different irradiation conditions as fixed parameters and the donors as random factors.

### Conditioned medium of senescent pellets had no major effect on cartilage formation

Senescent cells can affect their surrounding cells via the secretion of a SASP ^28^. To investigate whether or not the SASP contributes to reduced cartilage formation in chondrogenic pellets, conditioned medium of control and senescent pellets during chondrogenic differentiation (day 5-6 and day 12-13) was generated and added to non-irradiated recipient chondrogenic pellets at day 7 or day 14 (**Figure 4A**). First, we confirmed an increased expression of selected SASP factors *IL6* (*P*<0.001) and *MMP3* (*P*<0.001) in the irradiated pellets compared to non-irradiated control pellets (**Figure 4B**). Next, after exposition to the conditioned media of senescent pellets,, we observed that *COL2A1, ACAN, SOX9* and *COL1A1* expression were not significantly different compared to the control pellets cultured in control conditioned media (**Figure 4C**), suggesting that factors secreted from senescent cells during chondrogenesis do not directly alter the expression of chondrogenic genes in recipient pellets in our experimental conditions. To understand whether the absence of changes in the expression of chondrogenic markers was influenced by an altered expression in catabolic genes, we analyzed the expression of *MMP13, MMP1, MMP3* and *ADAMTS4*. Pellets cultured in conditioned medium of irradiated pellets had similar expression of catabolic genes as pellets cultured in control conditioned medium at both day-9 and day-16 of chondrogenic differentiation (**Figure 4C**). Overall, these results suggest that the SASP-factors produced by senescent cells in the pellets have no major effect at different stages of cartilage formation.

**Figure 4.**
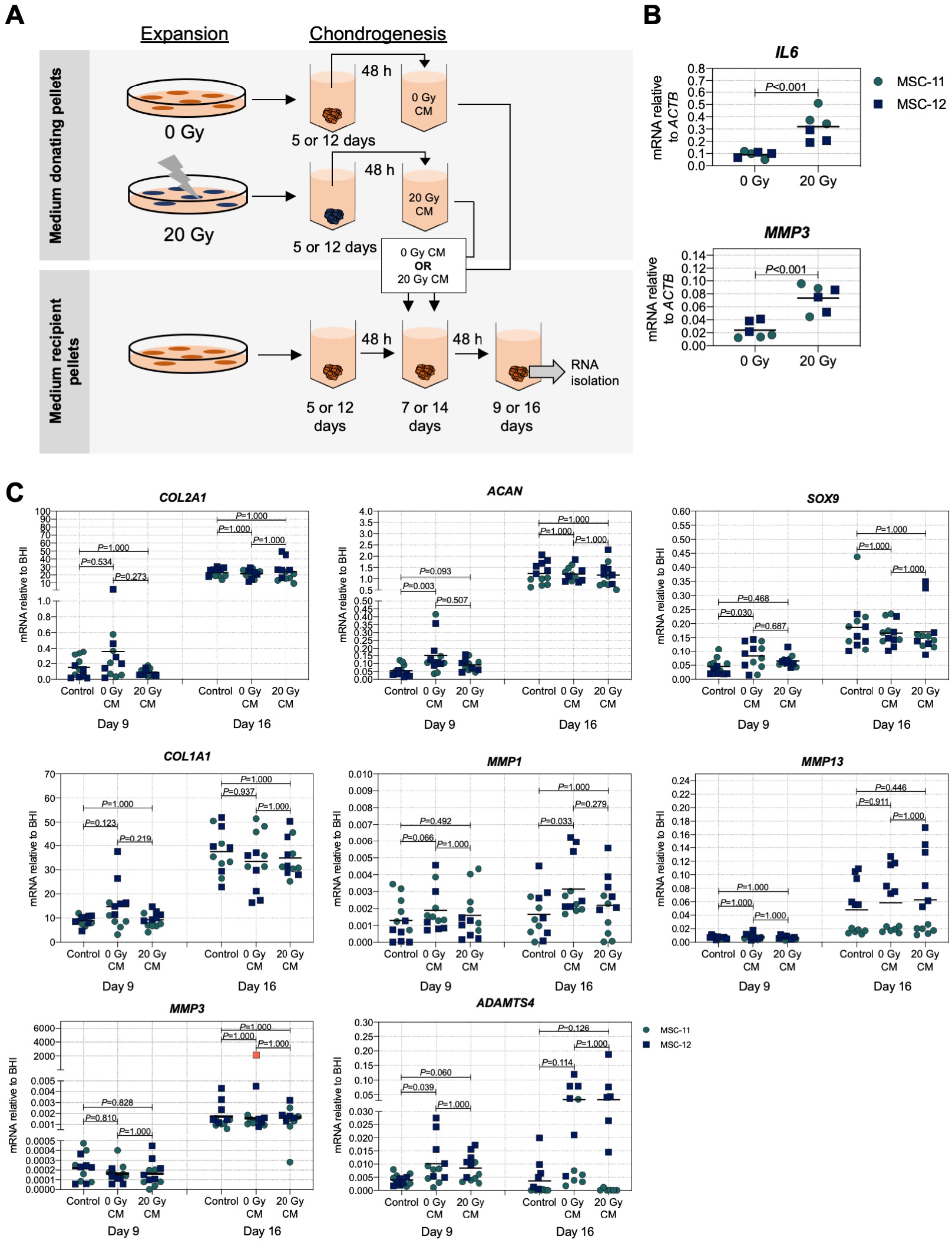
Conditioned medium from pellets of senescent MSCs did not alter the expression of chondrogenic genes in recipient non-senescent pellets. (A) Schematic overview of experimental setup. (B) mRNA expression of day 16 of chondrogenically differentiated pellets from 20 Gy gamma irradiated (during monolayer expansion) or not irradiated MSCs. N=2 donors with 3 replicates per donor. (C) mRNA expression of MSC pellets that were chondrogenically differentiated for 9 or 16 days and treated with conditioned medium for the last 48 h. The conditioned medium was obtained from chondrogenic pellets from MSCs that were irradiated with 20 Gy or not irradiated during expansion. Gene expression levels were normalized using best housekeeping index (BHI; *GADPH, HPRT* and *RSP27A*). MMP3 data shows one outlier in red.This value was excluded from the statistical analysis. N=2 donors with 6 replicates per donor. Data show individual data points and grand mean. *P*-values were obtained with the linear mixed model, using the different irradiation conditions as fixed parameters and the donors as random factors.

### The number of senescent cells is associated with a reduced cartilage production

Next, we asked if the observed negative effect on chondrogenesis was dependent on the number of senescent cells present at the moment of pellet formation. To answer these questions, we generated pellets starting with a different ratio of irradiated and non-irradiated cells and we monitored their chondrogenic differentiation capacity. The number of irradiation-induced senescent cells prior to chondrogenic differentiation was indeed associated with a reduced GAG and COL2 deposition (**Figure 5A-B** and **Supplementary Figure 4**) and both GAG and DNA content in chondrogenic pellets were negatively associated with the number of senescent MSCs (**Supplementary Figure 4B-C**). MSC pellets with 20-30% senescent MSCs had an average of 42% lower GAG content than pellets with non-irradiated cells (**Supplementary Figure 4**; *P*=0.008), suggesting that a low percentage of senescent cells already have a significant effect on the GAG deposition. MSC pellets with 45-55% senescent MSCs had, on average, 55% lower GAG/DNA than pellets with non-irradiated cells (**Supplementary Figure 4**; *P*=0.003), indicating that the non-senescent MSCs were still able to deposit GAG in the mixed pellets. Histological analysis showed clearly reduced GAG and COL2 deposition in MSC pellets with 45-55% senescent MSCs compared to pellets with non-irradiated cells (**Figure 5A-B** and **Supplementary Figure 5**). Furthermore, MSC pellets with 45-55% senescent MSCs had lower expression of *COL2A1, SOX9* and *ACAN* at day 21 of chondrogenic differentiation compared to non-senescent control MSCs, albeit not statistically significant (**Figure 5C**). MSC pellets with 70-80% senescent MSCs had a lower GAG content compared to MSC pellets with 45-55% senescent MSCs, however these pellets still deposited GAG (**Supplementary Figure 4**). On the other side, pellets with more than 90% senescent cells did no deposit GAG (**Figure 5A-B**). These pellets also had a low expression of *COL2A1* (97% reduced compared to non-senescent control MSCs; *P*=0.013), *SOX9* (75% reduced compared to non-senescent control MSCs, p=0.002) and *ACAN* (97% reduced compared to non-senescent control MSCs, p=0.014). No significant differences in the expression of *COL1A1* and the catabolic markers *MMP13, MMP1, MMP3, ADAMTS4* were observed between the different conditions (**Figure 5D-E**). These data may suggest that there is an inverse association between the number of senescent cells and the ability of generating cartilage.

**Figure 5.**
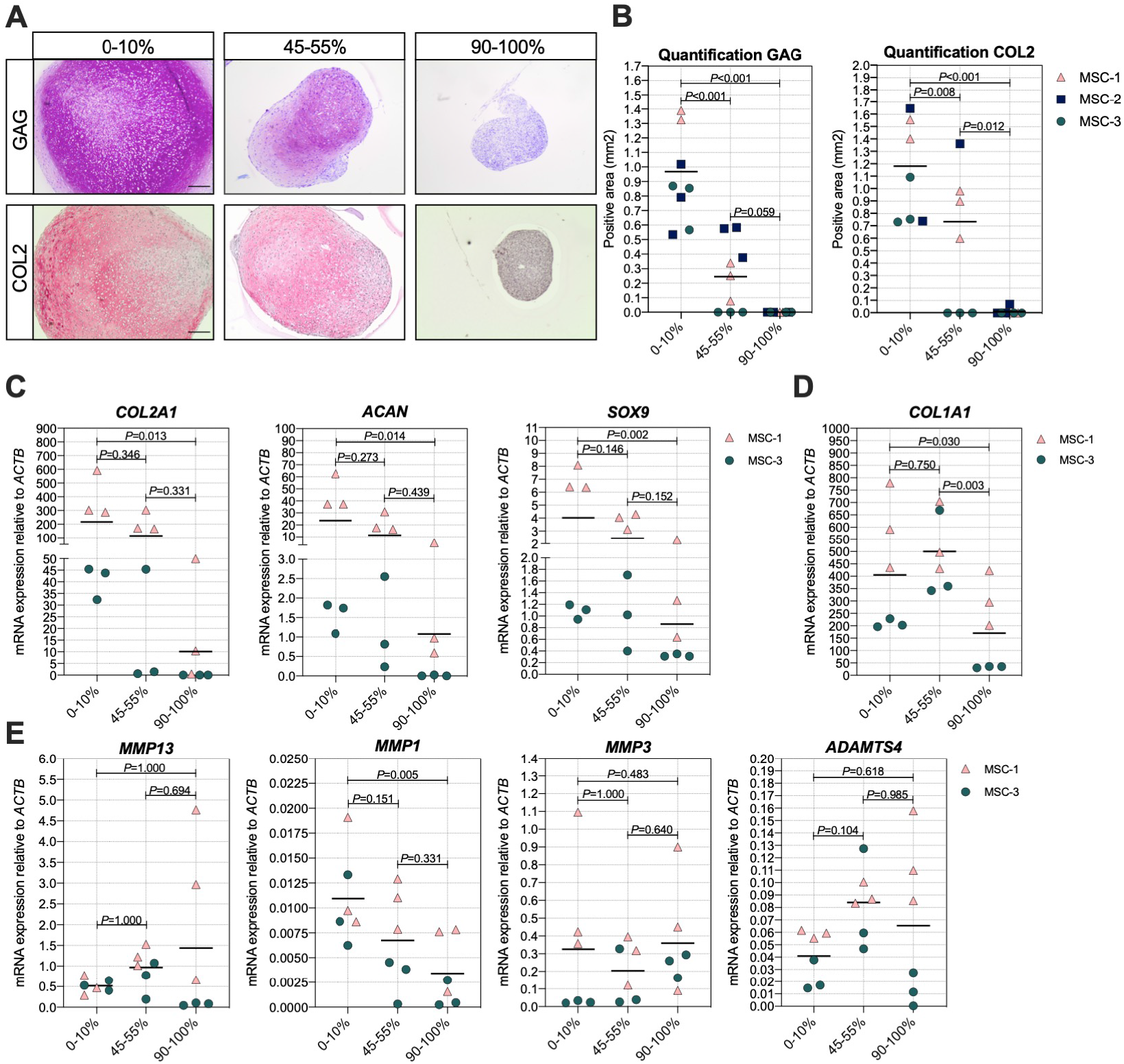
Higher ratio of senescent to non-senescent MSC resulted in less cartilage markers. (A-B) Representative images of Thionine and Collagen type 2 staining of MSCs that were gamma irradiated during expansion with 0 or 20 Gy, mixed and subsequently chondrogenically differentiated for 21 days. Scale bar represents 200 μm. N=3 donor with 2-3 pellets per donor. (C-E) mRNA expression of MSC pellets that were gamma irradiated during expansion with 0 or 20 Gy, mixed and subsequently chondrogenically differentiated for 21 days. Gene expression levels were normalized using *ACTB*. Data show individual data points and grand mean. *P*-values were obtained with the linear mixed model, using the experimental conditions as fixed parameters and the donors as random factors.

### Senescent MSCs are less responsive to TGFβ signaling

TGFβ is the main driver of chondrogenesis in MSC. In order to understand the reason why senescent cells have a reduced capacity to differentiate towards the chondrogenic lineage, we analyzed the TGFβ signaling activation by detecting the pSMAD2 levels in both irradiated MSCs (20 Gy) and control MSCs (0 Gy) upon TGFβ1 stimulation. In the presence of TGFβ1, pSMAD2 levels were higher in non-irradiated control MSCs compared to irradiated MSCs (**Figure 6A;** +TGFβ and **Figure 6B;** 6.9-fold, p=0.020), while no detectable pSMAD2 levels were present in MSCs without TGFβ1 stimulation (**Figure 6A**; -TGFβ). These data suggest that senescent MSCs are less responsive to TGFβ1, indicating that the reduced chondrogenic potential may be caused by a cell-intrinsic mechanism.

**Figure 6.**
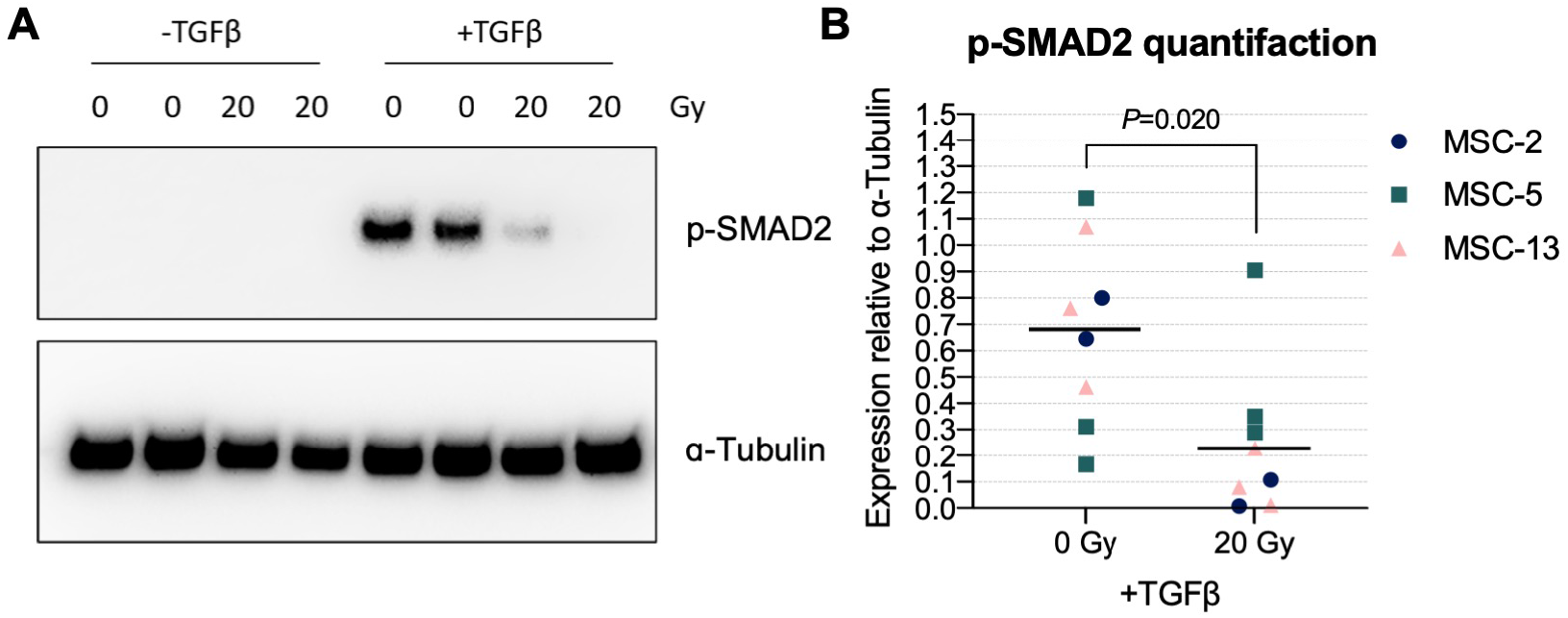
Senescent MSCs had low TGFβ induced phosphorylated SMAD2 levels. (A) Western blot for phosphorylated SMAD2 (p-SMAD2). (B) Quantification of western blot results relative to α-Tubulin. N=3 donors with 2-3 biological replicates per donor. Data show individual data points and grand mean. *P*-values were obtained with the linear mixed model, using the different experimental conditions as fixed parameters and the donors as random factors.

## Discussion

MSCs are promising cells for cartilage tissue regeneration therapies. To obtain reproducible and safe clinical outcomes it is necessary to understand how the chondrogenic differentiation capacity in MSC populations is regulated. In this study, we demonstrated that cellular senescence impairs the chondrogenic differentiation capacity of MSCs, we showed there is an association between the number of senescent cells at the start of the culture and the reduced chondrogenic differentiation potential, and we observed that senescent cells have a reduced ability to respond to TGFβ, the main factor responsible for chondrogenic differentiation of MSCs.

MSCs are a heterogeneous population of cells and the number of senescent cells varies between MSC cultures from different patients ^29^ and, most importantly, with passaging *in vitro* ^6, 18^. Here, we show for the first time that an increased number of senescent cells contribute to a reduced chondrogenic differentiation potential, indicating that the appearance of cellular senescence can contribute to heterogeneity in chondrogenic differentiation between MSC populations. This may be also linked with our previous observation that different MSC subtypes have a distinct differentiation capacity ^30, 31^. Furthermore, we show that in a mixed population with senescent MSCs, non-senescent MSCs are still able to differentiate towards the chondrogenic lineage and that the secretome of the senescent cells do not grossly influence the differentiation of neighboring non-senescent cells.

Our results suggest that senescent MSCs, while losing their chondrogenic differentiation potential, generally keep their osteogenic and adipogenic differentiation capacity. However, we identified some differences between gene expression and staining, specifically for the osteogenic assay. In fact, while mineral deposition seems slightly increased in irradiated MSCs, gene expression levels for osteogenic markers remain unaffected. This may explain why in the literature there is still no uniformed consensus on the effect of senescence in MSCs, with authors claiming minimal effect on osteogenic differentiation in late-passaged cells ^6^, others claiming upregulation ^32^ or even down-regulation of osteogenic differentiation ^33, 34^ with passaging or senescence. This discrepancy might also be linked with the timing of senescence induction during the experiments or could possibly be due to the different ways to induce senescence. Indeed, we and others previously observed different senescence phenotypes depending on the way senescence was induced ^25, 35^, and we cannot exclude that this may have a different impact on MSC differentiation.

In this study we demonstrated that the effect of irradiation-induced cellular senescence is largest during the early phases of chondrogenic differentiation. It has been shown that proliferation during the early phase of chondrogenesis is essential for proper chondrogenic differentiation ^36^. This indicates that impaired proliferation could be an explanation why MSCs failed to differentiate towards the chondrogenic lineage specifically when senescence is induced in monolayer or early during differentiation. Another explanation could be related to the differences we observed in the TGFβ signaling pathway activation in senescent MSCs compared to non-senescent MSCs. The TGFβ signaling has an important role in cartilage development and cartilage homeostasis ^37^. Particularly in the early phases of (re)differentiation, Smad2/3 phosphorylation is essential for chondrogenesis of MSCs ^38^ and for re-differentiation of de-differentiated chondrocytes ^39^. Here, we demonstrated that senescent MSCs have reduced pSMAD2 levels after TGFβ1 stimulation, compared to non-senescent control MSCs, suggesting that the canonical TGFβ signaling is altered in senescent MSCs. However, other non-canonical TGFβ pathways may be also involved in the process of cellular senescence and need further investigations.

It is known that senescent cells can affect the surrounding cells and tissues via their secretome. Previously, it has been shown that implantation of senescent cells can contribute to an OA-like phenotype in mice ^17^. In order to safely use MSCs for cartilage repair strategies, it is crucial to understand whether the SASP factors released by senescence cells can limit chondrogenesis or even contribute to cartilage degeneration. In this study, we found that the conditioned medium of chondrogenic pellets of senescent MSCs had no direct effect on the expression of the chondrogenic (*COL2A1, ACAN* and *SOX9)* or the catabolic (*MMP1, MMP13, MMP3* or *ADAMTS4)* markers in recipient pellet cultures. These data indicate that in our *in vitro* model, the SASP factors released from senescent MSCs have no negative effect on MSC chondrogenesis nor on the matrix degradation processes. Despite the absence of a direct effect on the MSCs exposed to the medium of senescent MSCs, we did find that senescent MSCs in the pellets had higher expression levels of inflammatory factors *IL6* and *MMP3*. The role of IL6 in cartilage tissue is controversial, since it has been shown to stimulate both cartilage degeneration and synthesis ^40-42^, and MMP3 promotes cartilage loss via degradation of multiple extracellular matrix components ^43^, indicating that the SASP factors released by MSCs could thus also contribute to the pathophysiology of OA. On the other hand, the SASP factors have been shown to be essential for tissue regeneration via the recruitment of macrophages ^44^. Therefore, more studies specifically focused on the role of individual SASP factors are necessary to better understand their role in cartilage generation and degeneration as well as possible interventions to counteract these effects.

In this study we explored how senescence in MSCs affect the chondrogenesis process. We showed that the number of senescent cells in MSC cultures is associated with a reduced chondrogenic differentiation potential. Especially senescence in the early phase of chondrogenesis could be detrimental for MSC-based cartilage tissue engineering. Therefore strategies that prevent or abolish senescence in MSCs could be beneficial for MSC-based cartilage repair.

## Author contributions

Study conception, design, analysis and final approval of the article: Chantal Voskamp, Wendy J. L. M. Koevoet, Gerjo J.V.M. van Osch, Roberto Narcisi. Drafting the article: Chantal Voskamp, Gerjo J.V.M. van Osch, Roberto Narcisi.

