## Supplementary information for "Senescence during early differentiation reduced chondrogenic differentiation capacity of mesenchymal progenitor cells"

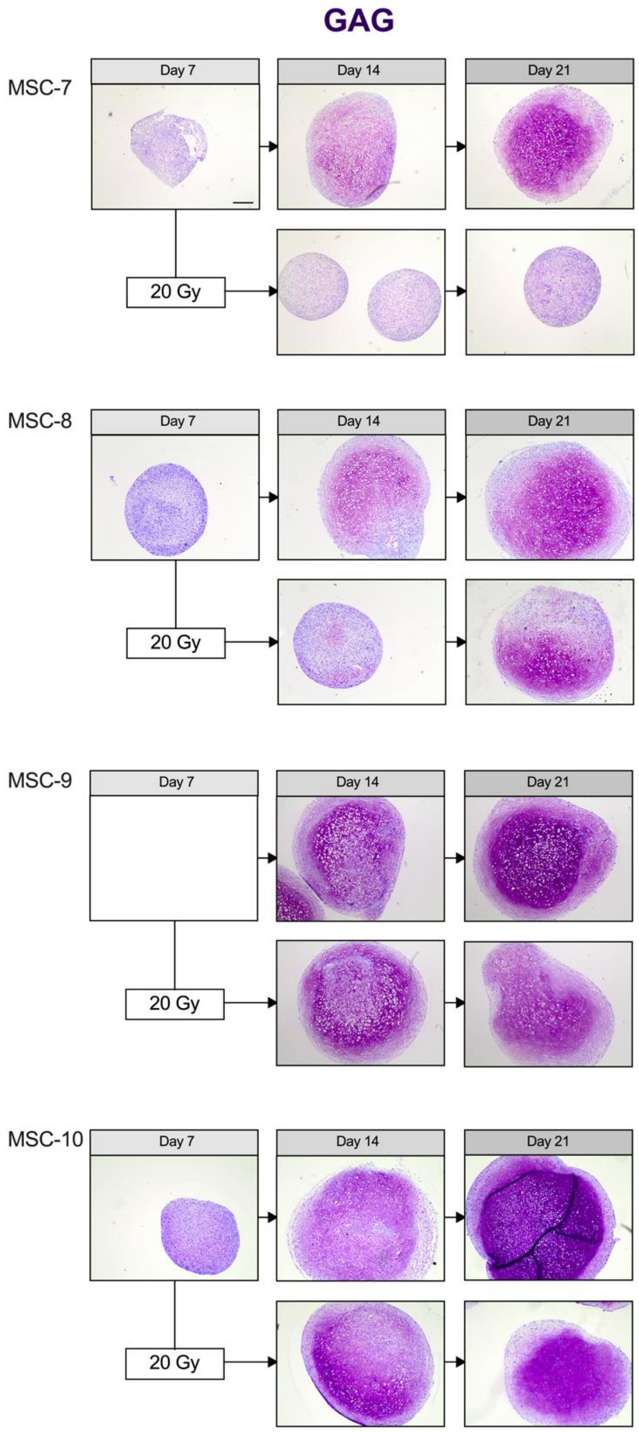


**Supplementary Figure 1 –Thionine staining of irradiated MSC pellets at day 7.** Images of Thionine (GAG) staining of MSC control pellets that were chondrogenically differentiated for 7, 14 and 21 days or MSC pellets that were irradiated at day 7 during chondrogenic differentiation and subsequently differentiated for 7 or 14 days. The day 7 pellets of donor MSC-9 are missing due to a technical issue during processing. The scale is the same in all images. Scale bar represents 200 µm and is indicated in the day 7 pellet of donor MSC-7. The images of donor MSC-8 are the same as depicted in Figure 2A. N=4 donors with 2-3 pellets per donor.

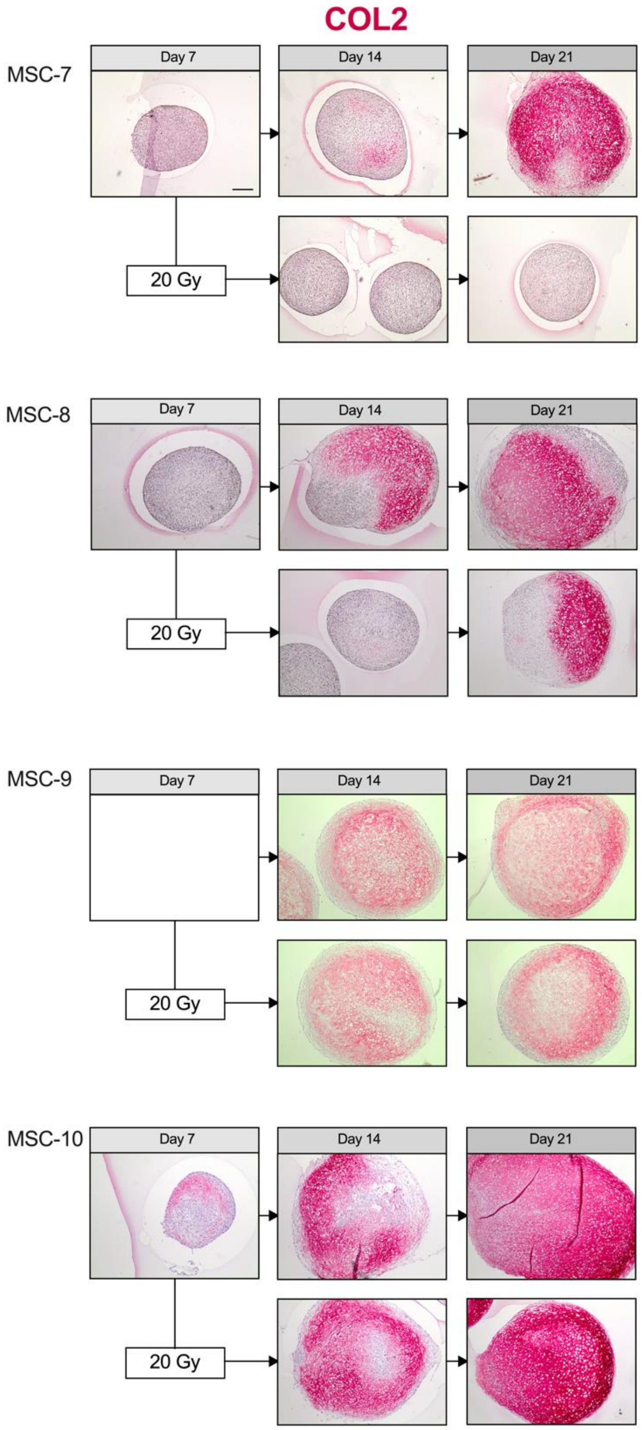


**Supplementary Figure 2 – Collagen type 2 staining of irradiated MSC pellets at day 7.** Images of Collagen type 2 (COL2) immunohistochemical staining of MSC control pellets that were chondrogenically differentiated for 7, 14 and 21 days or MSC pellets that were irradiated at day 7 during chondrogenic differentiation and subsequently differentiated for 7 or 14 days. Positive staining in red. The day 7 pellets of donor MSC-9 are missing due to a technical issue during processing. The scale is the same in all images. Scale bar represents 200 µm and is indicated in the day 7 pellet of donor MSC-7. The images of donor MSC-8 are the same as depicted in Figure 2B. N=4 donors with 2-3 pellets per donor.

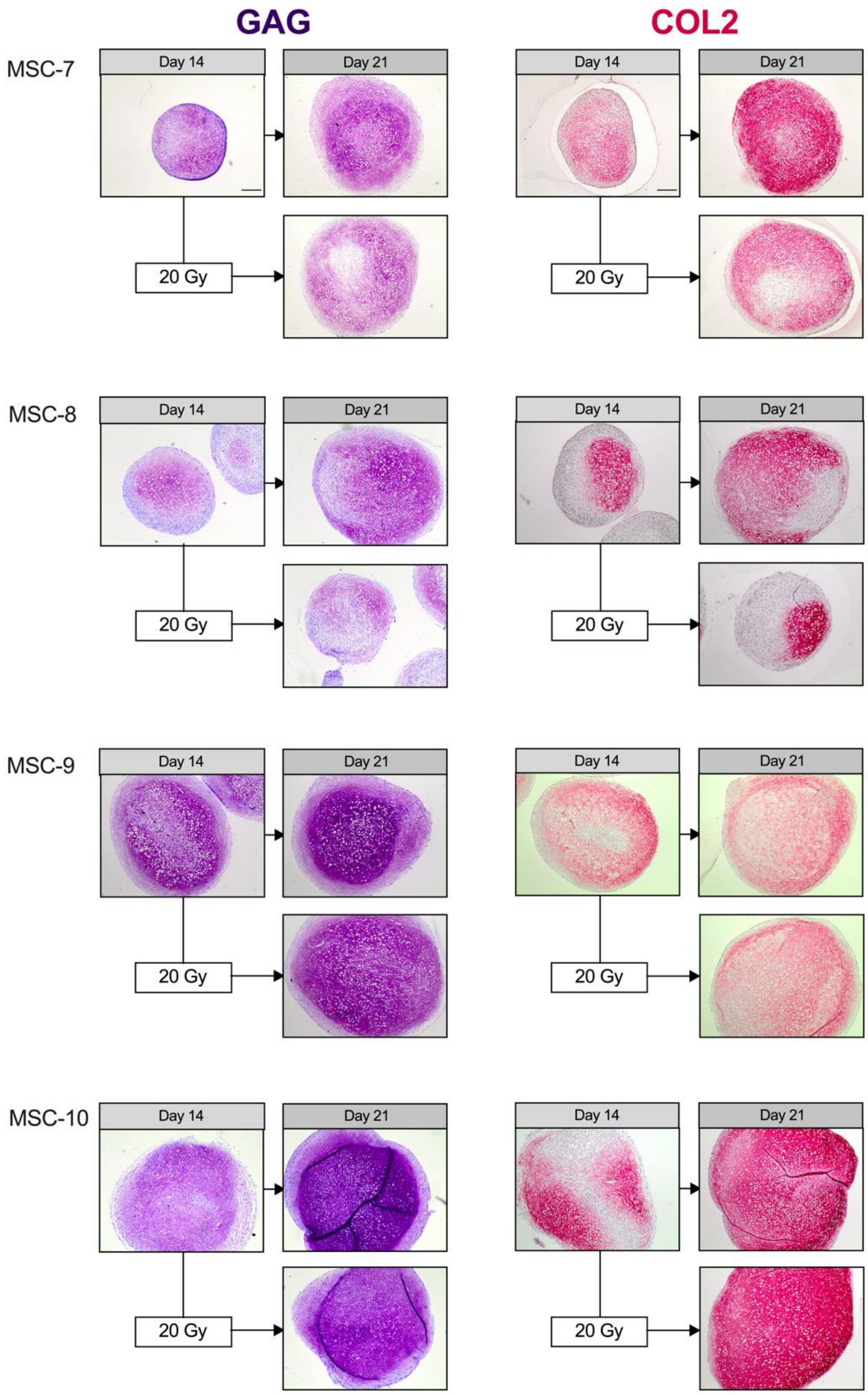


**Supplementary Figure 3 – Thionine and Collagen type 2 staining of irradiated MSC pellets at day 14.** (Left panels) Images of Thionine (GAG) and (right panels) images of Collagen type 2 (COL2) staining of MSC control pellets that were chondrogenically differentiated for 14 and 21 days or MSC pellets that were irradiated at day 14 during chondrogenic differentiation and subsequently differentiated for 7. The scale bar represents 200 µm. The images of donor MSC-7 are the same as depicted in Figure 3A-B. N=4 donors with 2-3 pellets per donor.

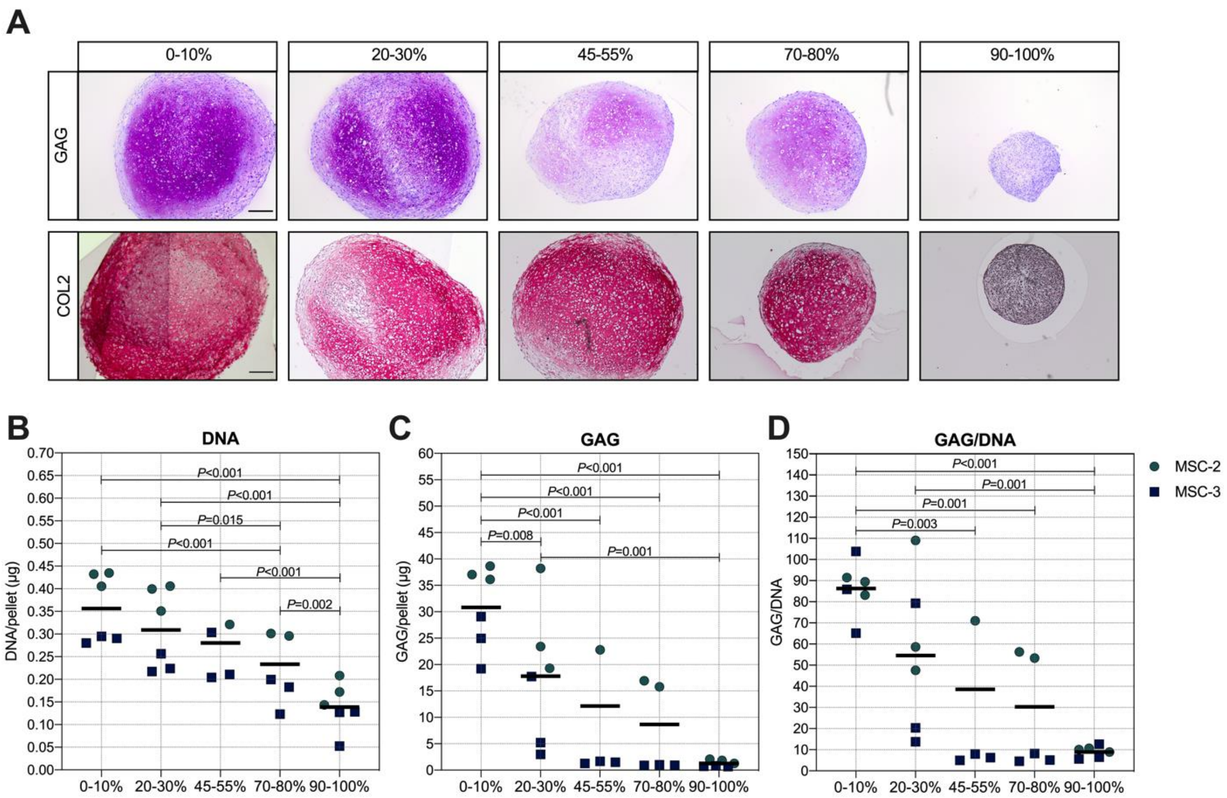


**Supplementary Figure 4 - GAG and DNA content in MSCs pellets with senescent and non-senescent cells mixed.** (A) Representative images of Thionine (GAG) and Collagen type-2 (COL2) staining of MSCs that were gamma irradiated during expansion with 0 or 20 Gy, mixed (percentages indicate the percentage of senescent MSCs) and subsequently chondrogenically differentiated for 21 days. Scale bar represents 200 µm. N=2 donors with 2-3 pellets per donor. (B-D) GAG, DNA and GAG/DNA content of MSCs that were gamma irradiated during expansion with 0 or 20 Gy, mixed (percentages indicate the percentage of senescent MSCs) and subsequently chondrogenically differentiated for 21 days. N=2 donors with 2-3 pellets per donor. *P*-values were obtained with the linear mixed model, using the different experimental conditions as fixed parameters and the donors as random factors and Bonferroni post-hoc test was used to correct for multiple comparisons.

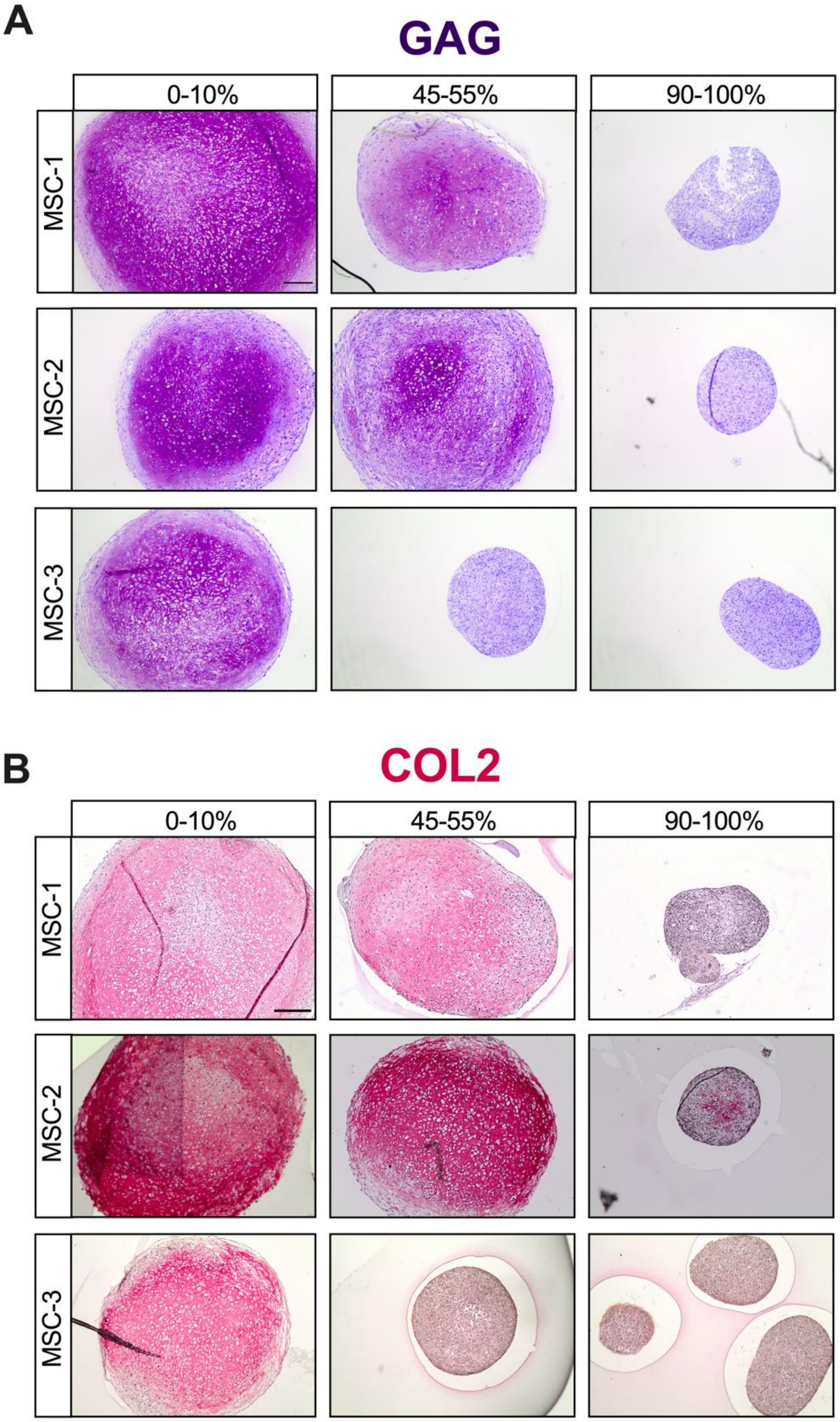


**Supplementary Figure 5 – Thionine and Collagen type 2 staining of MSC pellets with different ratios of senescent MSCs.** (A) Thionine and (B) Collagen type 2 staining of MSCs that were gamma irradiated during expansion with 0 or 20 Gy, mixed and subsequently chondrogenically differentiated for 21 days. Representative images from different technical triplicates are depicted Scale bar represents 200 µm. N=3 donors with 2-3 pellets per donor. The images of donor MSC-1 are the same as depicted in Figure 5A. The images of the Collagen type 2 staining with 0-10% and 90-100% senescent MSCs for donor MSC-1 and MSC-3 are the same as depicted in Figure 1D.
